# Insights on early mutational events in SARS-CoV-2 virus reveal founder effects across geographical regions

**DOI:** 10.1101/2020.04.09.034462

**Authors:** Carlos Farkas, Francisco Fuentes-Villalobos, José Luis Garrido, Jody J Haigh, María Inés Barría

## Abstract

Here we aim to describe early mutational events across samples from publicly available SARS-CoV-2 sequences from the sequence read archive repository. Up until March 27, 2020, we downloaded 53 illumina datasets, mostly from China, USA (Washington DC) and Australia (Victoria). Of 30 high quality datasets, 27 datasets (90%) contain at least a single founder mutation and most of the variants are missense (over 63%). Five-point mutations with clonal (founder) effect were found in USA sequencing samples. Sequencing samples from USA in GenBank present this signature with 50% allele frequencies among samples. Australian mutation signatures were more diverse than USA samples, but still, clonal events were found in those samples. Mutations in the helicase and orf1a coding regions from SARS-CoV-2 were predominant, among others, suggesting that these proteins are prone to evolve by natural selection. Finally, we firmly urge that primer sets for diagnosis be carefully designed, since rapidly occurring variants would affect the performance of the reverse transcribed quantitative PCR (RT-qPCR) based viral testing.

## Introduction

The COVID-19 pandemic caused by a novel 2019 SARS coronavirus, known as SARS-CoV-2, is rapidly evolving worldwide, surpassing the 8000 total cases of 2002-2004 SARS coronavirus outbreak (SARS-CoV-1) after one month of the initially identified case on December 31, 2019, in Wuhan city, China (Wilder-Smith et al. 2020). As SARS-CoV-2 is human-to-human transmitted, it is a threat to the global population. It is critical to understand SARS-CoV-2 characteristics to deal with this ongoing pandemic and to develop future treatments. SARS-CoV-2 virus is an enveloped, positive-stranded RNA virus with a large genome (29.9 kb) belonging to the family Coronaviridae, order Nidovirales (de Wit et al. 2016). One of the striking genomic features of this novel virus is the presence of a novel furin-like cleavage site in the S-protein of the virus, which differs from SARS-CoV-1 and may have implications for the life cycle and pathogenicity of the novel virus (Coutard et al. 2020; Wu et al. 2020a). Firstly, it was suggested that SARS-CoV-2 is a close relative of the RaTG13 bat-derived coronavirus (around 88% identity) rather than of SARS-CoV-1 (79% identity) or middle east respiratory syndrome coronavirus MERS-CoV (50% identity) (Lu et al. 2020). Due to this association with bat coronaviruses, it was also argued that SARS-CoV-2 virus has the potential to spread into another species, as bat coronaviruses do (Hu et al. 2018). Recently, it was demonstrated that SARS-CoV-2 is closely related to a pangolin coronavirus (Pangolin-CoV) found in dead Malayan pangolins with a 91.02% identity, the closest relationship found so far for SARS-CoV-2 (Zhang et al. 2020). In that study, genomic analyses revealed that the S1 protein of Pangolin-CoV is related closer to SARS-CoV-2 than to RaTG13 coronavirus. Also, five key amino acid residues involved in the interaction with the human ACE2 receptor are maintained in Pangolin-CoV and SARS-CoV-2, but not in RaTG13 coronavirus. Thus, it is likely that pangolin species are a natural reservoir of SARS-CoV-2-like coronaviruses and SARS-CoV-2 will continue to evolve with novel mutations, as the pandemic evolves. In this scenario, it is expected that diverse signatures of viral variants spread among different populations in the world. Recently, thousands of GenBank sequences from SARS-CoV-19 available at the NCBI virus database were trackable by region, suggesting that the transmission occurred mainly through clonal events due to clustering of the available sequences (Chen et al. 2020; Kupferschmidt 2020) (https://www.ncbi.nlm.nih.gov/labs/virus/vssi/#/virus?SeqType_s=Protein&VirusLineage_ss=SARS-CoV-2,%20taxid:2697049). As a proof of concept, in the early beginning of the outbreak in China, sequencing the virus from nine patients from Wuhan in China revealed 99.9% similarity among samples. That finding suggests 2019-nCoV originated from one source within a very short time, supporting clonality of spreading (Lu et al. 2020). In this study, we characterized the early mutational events across 30 high-quality datasets publicly available on the sequence read archive repository. 27 out of 30 samples (90%) contain at least a single founder mutation and most of the variants across samples are missense (over 63%). Remarkably, SARS-CoV-2 variants in USA samples display a clonal pattern of acquired mutations, that are dissimilar to Australian SARS-CoV-2 mutations, which were found to be heterogeneous. A mutational signature from USA mutations was however found in an Australian sample, suggesting a worldwide spread of this molecular signature consisting of five-point mutations. Remarkably, mutations in the helicase and orf1a proteins of the virus were found more frequently than others, suggesting that these proteins are prone to evolve throughout natural selection. As proof of the latter, a single nucleotide polymorphism (SNP) in an Australian sample causes a bona-fide stop codon in the helicase protein, strongly suggest this protein will evolve on SARS-CoV-2 in the future. As genetic drift prompts the mutational spectrum of the virus, we recommend frequently sequencing the viral pool in every country to detect the founder events relevant for SARS-CoV-2 testing in each population

## Materials & Methods

### Data Collection

Raw illumina sequencing data were downloaded from the following NCBI SRA BioProjects: SRA: PRJNA601736 (Chinese datasets), SRA: PRJNA603194 (Chinese dataset) (Wu et al. 2020b), SRA: PRJNA605907 (Chinese datasets) (Shen et al. 2020), SRA: PRJNA607948 (USA-Wisconsin datasets), SRA: PRJNA608651 (Nepal dataset), SRA: PRJNA610428 (USA-Washington datasets), SRA: PRJNA612578 (USA-San-Diego dataset), SRA: PRJNA231221 (USA-Washington dataset) (Sichtig et al. 2019), SRA: PRJNA613958 (Australian-Victoria datasets), SRA: PRJNA231221 (USA-Maryland dataset), and SRA: PRJNA614995 (USA-Utah datasets). All SRA accessions are depicted in Supplementary Table 1, sheet 1.

### Data processing

Raw reads were aligned with bowtie2 aligner (v2.2.6) (Langmead & Salzberg 2012) against SARS-CoV-2 reference genome NC_045512.2 (https://www.ncbi.nlm.nih.gov/nuccore/NC_045512), using the following parameters: -D 20 -R 3 -N 0 -L 20 -i S,1,0.50. Samtools v1.9 (using htslib v1.9) (Li et al. 2009) was used to sort sam files, remove duplicate reads and index bam files. bcftools v1.9 (part of the samtools framework) was used to obtain depth of coverage in each aligned sample. For variant calling from each sample, bcftools mpileup was used with the following parameters: -B -C 50 -d 250. To obtain founder mutations, strict filtering of called variants was performed with bcftools filter, considering variants only with Mann-Whitney U test of read position bias over 0.1 and the number of high-quality reference alleles divided by high-quality alternate alleles over 0.3. All commands to obtain these computational steps are publicly available at https://github.com/cfarkas/SARS-CoV-19_illumina_analysis.

### SNVs consequences and classification

All SNP and INDELs consequences were assessed in each sample by using snippy haploid variant calling and core genome alignment pipeline: https://github.com/tseemann/snippy.

### Construction of multiple sequence alignments with GenBank sequences

436 GenBank SAR2-CoV-2 submitted fasta sequences were downloaded on March 31, 2020, from NCBI virus database (https://www.ncbi.nlm.nih.gov/labs/virus/vssi/#/) using as query “Severe acute respiratory syndrome coronavirus 2 (SARS-CoV-2), taxid:2697049”. These sequences were aligned and processed with the same steps as we did with the fastq datasets. (see Data processing section and “Alignment of GenBank Sequences, March 31, 2020” in https://github.com/cfarkas/SARS-CoV-19_illumina_analysis).

### Primer List obtention

CDC primers currently in use (April 2020) were obtained from https://www.cdc.gov/coronavirus/2019-ncov/lab/rt-pcr-panel-primer-probes.html, generated by the Division of Viral Diseases, National Center for Immunization and Respiratory Diseases, Centers for Disease Control and Prevention, Atlanta, GA, USA. University of Hong-Kong primers currently in use from the were obtained from https://www.who.int/docs/default-source/coronaviruse/peiris-protocol-16-1-20.pdf?sfvrsn=af1aac73_4 and Institute Pasteur primers currently in use were obtained based on the first sequences of SARS-CoV-2 made available on the GISAID database (Shu & McCauley 2017) on January 11, 2020, available here: https://www.who.int/docs/default-source/coronaviruse/real-time-rt-pcr-assays-for-the-detection-of-sars-cov-2-institut-pasteur-paris.pdf?sfvrsn=3662fcb6_2. We also included literature-based primers (Kim et al. 2020) and ten primer-BLAST (Ye et al. 2012) designed primers against SARS-CoV-2 reference genome NC_045512.2.

## Results

### Inspection of variants reveals well-defined signatures with founder effect across sequenced samples

We downloaded 53 raw datasets from the sources referred in Material and Methods (see **Supplementary Table 1**, sheet 1) and we aligned fastq reads against SARS-CoV-2 reference genome NC_045512.2, corresponding to the initial isolate Wuhan-Hu-1. Out of 53 total datasets, only 30 displayed coverage over 30X, suitable for the variant calling framework, which we subsequently analyzed in depth. We also manually checked the coverage of each sample by using the Integrative Genomics Viewer (IGV) tool (Robinson et al. 2011). Variant calling reveals a diverse number of variants in each sample, which were strictly filtered to obtain founder mutations in each sample (see Material and Methods). Five datasets from Chinese origin were selected after coverage filtering and only two of them displayed founder mutations (see **Supplementary Table 1**, sheet 1). Fourteen datasets from the USA were filtered (USA-Washington study, hereafter referred as USA-WA) and remarkably, all of them displayed variants presenting a defined mutational signature, consisting in a core of five founder mutations at positions 8782, 17747, 17858, 18060 and 28144 of the SARS-CoV-2 reference genome (see **Figure 1A, Supplementary Table 1**, sheet 1). Mutational landscape of SARS-CoV-2 samples in Australia (Australia-Victoria samples, hereafter Australia-VIC) was clearly heterogeneous, displaying a variety of founder mutations per sample but also shared variants were observed within samples. One variant (position 26144) is present in 5/11 Australia-VIC samples and variants with positions 8782 and 28144 are present in 3/11 Australian samples (see **Figure 1A, Supplementary Table 1**, sheet 1). Notably, one Australian-VIC sample (SRR11397717) displayed the same five-point mutational signature of USA-WA samples, two samples contain the same mutational signature presenting one deletion (SRR11397715 and SRR11397716) and one novel signature (SRR11397728) presents an SNP that creates a stop codon (see **Figure 1A**, respectively). All of these called variants present mutant allele frequencies near or equal to 100%, evidenced in the number of mutant_alleles / reference_alleles (see mutant allele frequency in **Supplementary Table 1**, sheet 2 and 3, respectively) easily visualized in the aligned bam files (see **Figure 1B** from USA-WA and Australian-VIC samples, respectively). These analyses suggest these variants were already spread in the infected population in the early days of the outbreak, they are not restricted by country and that they will continue to spread along with the growing cases. To support the latter, we downloaded 436 GenBank sequences of SARS-CoV-2 available in NCBI virus database from different countries and we aligned them with the SARS-CoV-2 reference genome. Variant calling from these alignment reveals the substantial presence of these USA-WA signature, with allele frequencies ranging 37-46% (see **Figure 1B**, bottom and **Supplementary Table 1**, sheet 4). Thus, the USA-WA signature is widespread among SARS-CoV-2 infections likely due to founder effect. To figure out whether this signature is enriched in USA samples, we performed the same analysis in a subset of 277 sequences, all of them from USA origin. As expected, the USA-WA signature consisting of five-point mutations are enriched in allele frequencies over 50% plus (see **Supplementary Table 1**, sheet 5). Also, in these 277 sequences, six more variants were detected with smaller allele frequencies as reported for USA-WA variants. It is expected that these variants also will rise among infected people in the USA as the USA-WA signature did. A summary of the USA-WA mutational signature is depicted in Table 1.

**Figure 1:**
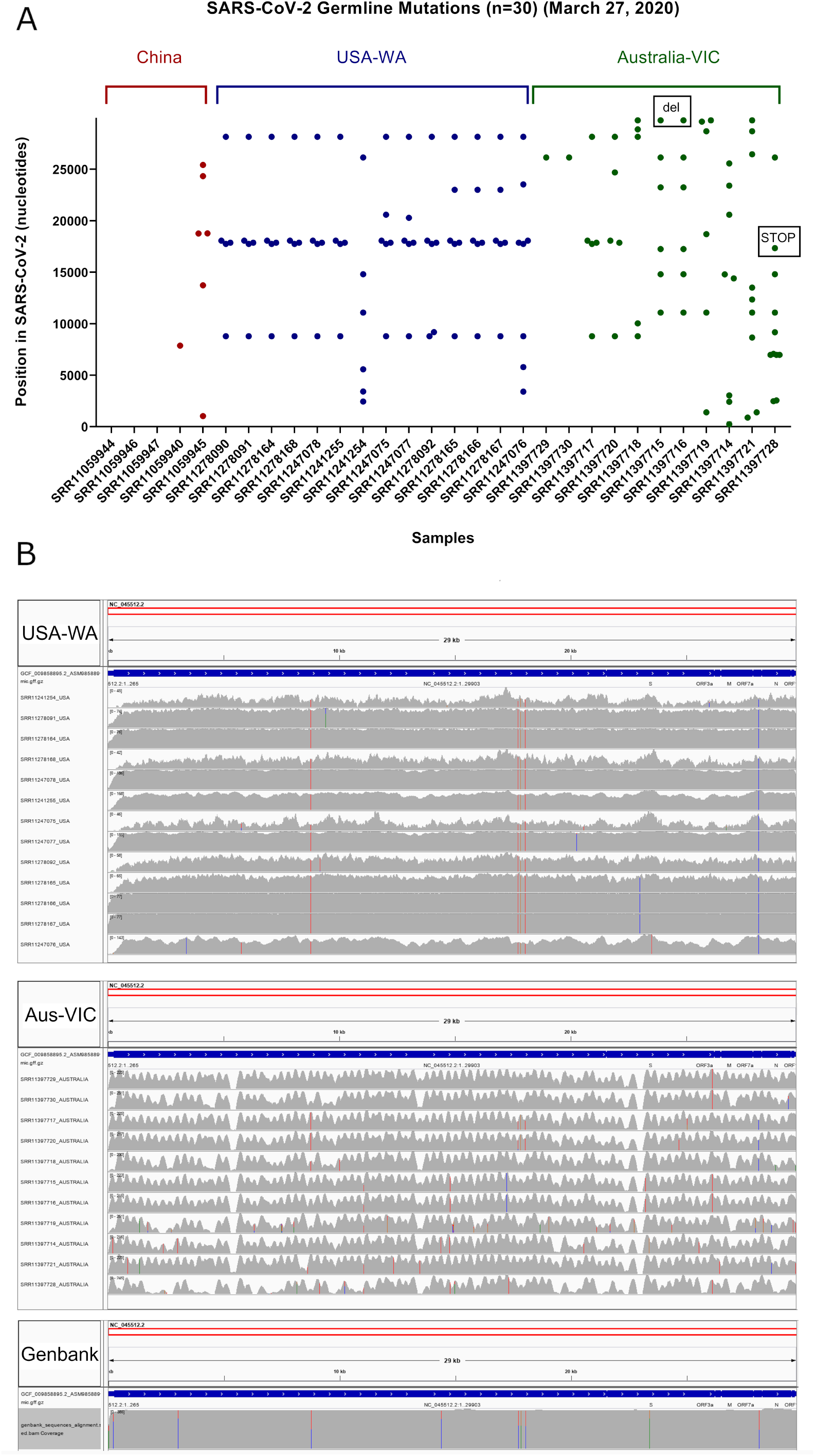
Inspection of variants reveals well-defined signatures with founder effect across sequenced samples. A) Plot of germline variants sorted by country (China=red, USA=blue and Australia=green) and by the number of variants, from left to right. Deletions and stop codons are framed with black rectangles. B) Top IGV screenshots of coverage from USA samples (n=13) aligned against SARS-CoV-2 reference genome. Germline variants are depicted with colored lines. Middle IGV screenshots of coverage from Australian samples (n=11) aligned against SARS-CoV-2 reference genome. Germline variants are depicted with colored lines. Bottom IGV screenshots of coverage from GenBank sequenced samples of SARS-CoV-2 (n=436) aligned against SARS-CoV-2 reference genome. Variants are depicted with colored lines.

**Table 1:** Allele frequencies of five detected variants in 436 genbank available sequences

| **POS** | **REF** | **ALT** | **INFO** | **AF** |
| --- | --- | --- | --- | --- |
| 8782 | C | T | DP=302; DP4=163,0,139,0;MQ=42 | 46.02649 |
| 17747 | C | T | DP=302; DP4=188,0,114,0;MQ=42 | 37.74834 |
| 17858 | A | G | DP=302; DP4=188,0,114,0;MQ=42 | 37.74834 |
| 18060 | C | T | DP=302; DP4=183,0,119,0;MQ=42 | 39.40397 |
| 28144 | T | C | DP=303; DP4=163,0,140,0;MQ=42 | 46.20462 |

POS: position in SARS-CoV-2 reference sequence NC\_045512.2, REF: reference allele, ALT: mutant allele, DP4: Number of high-quality ref-forward , ref-reverse, alt-forward and alt-reverse bases, AF: allele frequency.

### Classification of founder mutations

Classification of variants performed by snippy tool analysis reveals that most of the referred variants in USA-WA and Australia-VIC samples are preferentially missense (63% for USA-WA samples and 74% for Australia-VIC samples, respectively) rather than synonymous (see **Figure 2A, Supplementary Table 1**, sheets 2 and 3, respectively). Focusing on missense variants of USA-WA signature, the most recurrent mutations occurred in the helicase, 3’-to-5’ exonuclease and ORF8 protein, accounting for the 81% of missense variants in this signature (see **Figure 2B, Supplementary Table 1**, sheet 2). These mutations correspond to the five core point mutations observed in USA-WA samples. In the case of Australia-VIC missense samples, the scenario is more complex, due to the heterogeneity of the signatures. Mutations in the orf1a polyprotein, ORF3a protein, helicase and surface glycoprotein account for 69% of the missense variants present in the Australia-VIC signatures. Importantly, two Australian samples (SRR11397715 and SRR11397716) present the same mutational profiling with deletions in the stem loop of the virus and notably, one sample (SRR11397728) presents an SNP that created a stop codon in the helicase protein (see **Supplementary Table 1**, sheet 3). Since every USA-WA and Australia-VIC sample that presents variants in the helicase gene contains at least one missense variant, this evidence strongly suggests that this gene is under active evolution it will likely continue to evolve along with SARS-CoV-2 pandemic.

**Figure 2:**
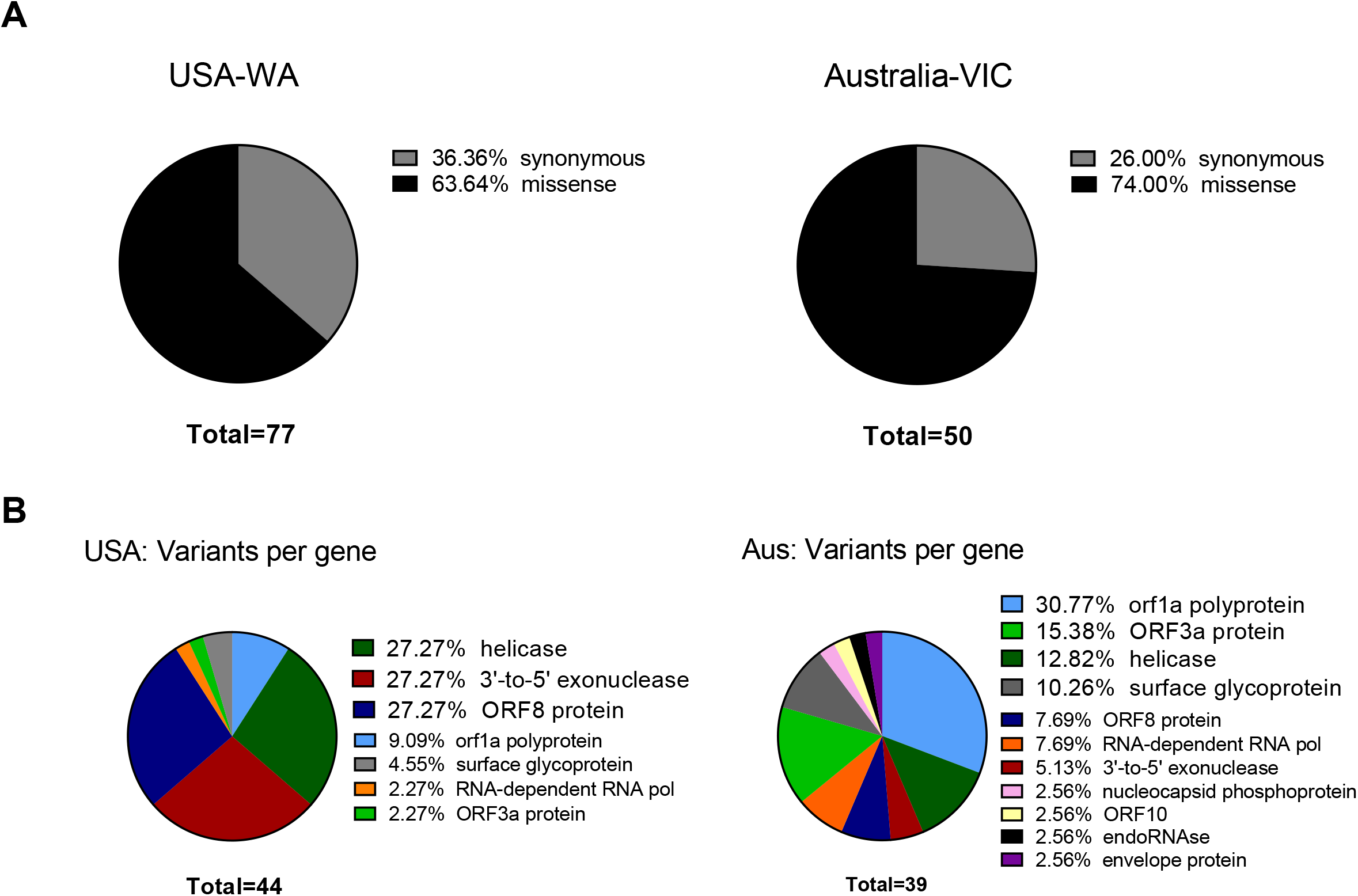
Classification of founder mutations. A) Left: Ratio of Synonymous / missense variants across USA samples. Missense variants are denoted in black and Synonymous variants are depicted in grey N indicates the total number of classified variants. Right: Same as left for Australian samples. B) Left: Missense variants classification from USA samples across SARS-CoV-2 genes. Each color indicates a different gene stated in legend (at right of the pie charts). Right: Same as left from Australian samples.

### SNPs in SARS-CoV-2 can diminish efficiency of RT-qPCR testing

We downloaded the Centers for Disease Control and Prevention primer list, consisting of three primer sets designed against ORF9 structural protein (nucleocapsid phosphoprotein), each one with a fluorescent probe for reverse transcriptase quantitative PCR (see https://www.cdc.gov/coronavirus/2019-ncov/lab/rt-pcr-panel-primer-probes.html and https://github.com/cfarkas/SARS-CoV-19_illumina_analysis, file CDC_primers.fasta). We investigated if these primers hybridized at positions with the variants reported herein from the 30 next-generation sequencing datasets. We aligned these primer sequences with SARS-CoV-2 reference genome and the 30 analyzed samples. We found two Australian clonal samples (SRR11397719 and SRR11397721) presenting one founder synonymous SNP (position 28688) that falls into the 2019_nCoV_N2_Forward_Primer hybridization region. In the first sample, we also found an SNP with low allele frequency that falls into the 2019-nCoV_N1_Reverse_Primer (see **Figure 3**). We also challenged a list of primers available from literature (16), primers currently used for the Institute Pasteur, Hong Kong University and ten primers obtained from primer-blast (see Material and Methods for sources) against 206 variants consisting in GenBank variants collected from USA samples including variants from the next-generation sequencing datasets employed here (see **Supplementary Table 1**, sheet 6). Of these, CDC primer set 2019-nCoV_N3 forward and reverse along with its probe were discarded again, due to potential reduced efficiency during priming. Similarly, sets 2, 3 and 8 from primer-blast, were discarded. Conversely, primers from Pasteur Institute and Hong-Kong university passed the filter, respectively. Thus, increasing variation in SARS-CoV-2 can confound quantitative RT-qPCR in the future depending on the primer design.

**Figure 3:**
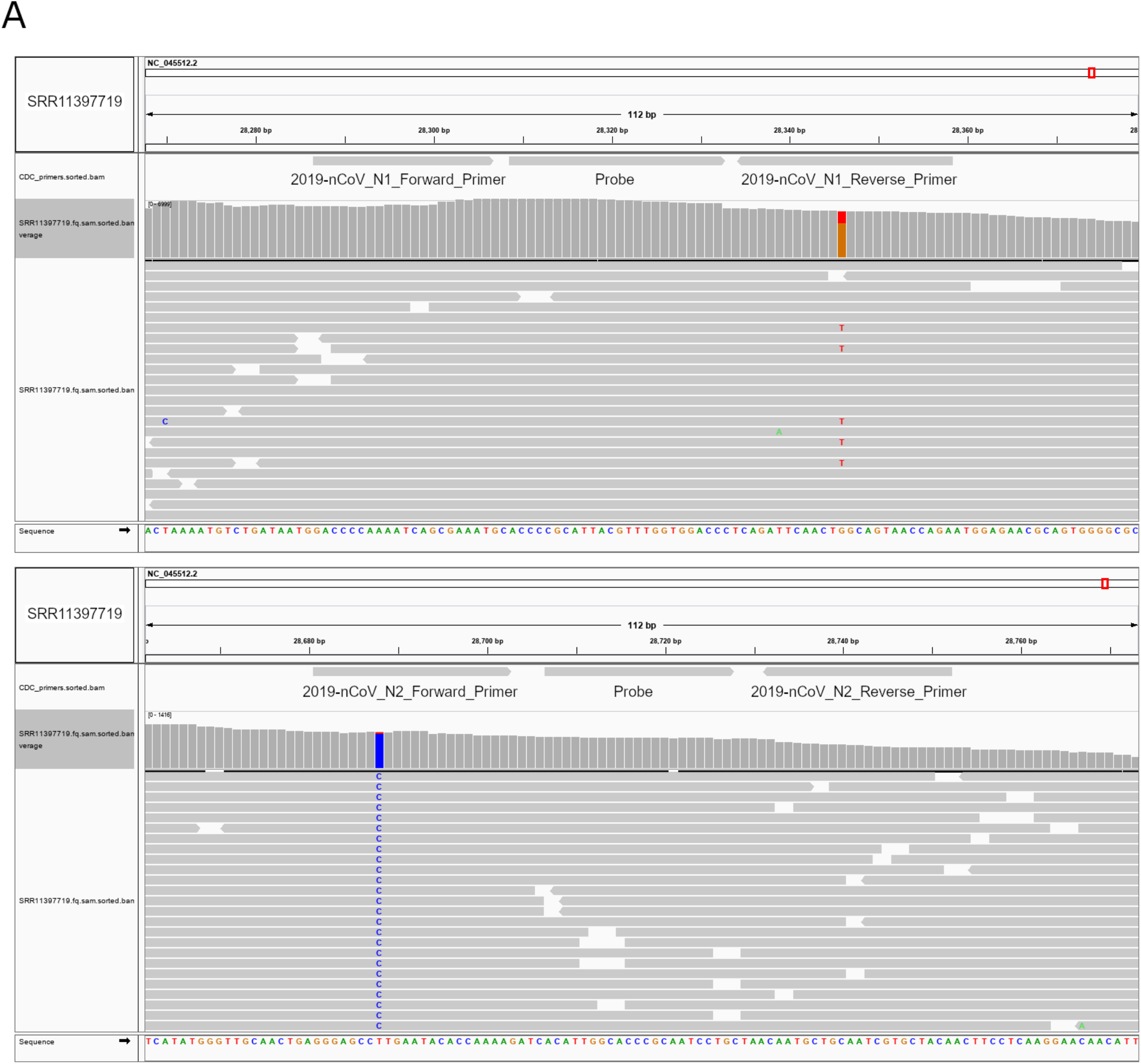
SNPs in SARS-CoV-2 can diminish efficiency of RT-qPCR testing. A) Top IGV screenshots of coverage from Australian sample SRR11397719 aligned against SARS-CoV-2 reference genome. A low allele frequency variant is depicted in red (T). Primers tracks are denoted at the top of the screenshot along with SARS-CoV-2 gff gene models. Bottom Same as Top for sample SRR11397719 denoting a germline variant in blue.

## Discussion

Since SARS-CoV-2 is an RNA virus that rapidly evolves after infection, these evolutionary events will likely affect its fitness over time. In this study, we reveal the early mutational events in small samples from mainly two geographically unrelated populations (Washington, USA, and Victoria, Australia) and we compared these results with variants observed in submitted GenBank sequences in NCBI viral portal, from different countries. As already reported with HIV and Chikungunya outbreaks, the founder effect of five-point mutations was observed in almost all USA-WA samples obtained by next-generation sequencing. These mutations also have a high allele frequency (around 50%) in SARS-CoV-2 GenBank sequences from USA origin (Foley et al. 2000; Tsetsarkin et al. 2011). These mutational signatures are likely to be overrepresented among Washington infections and USA infections overall, if not globally. Supporting this, the alignment of 435 submitted sequences to NCBI virus from different countries (March 31, 2020) shows allele frequencies of 37-46% of these five SNPs across samples worldwide. These SNPs cause missense mutations on helicase, 3’-to-5’ exonuclease and ORF8 proteins. In the case of Victoria samples from Australia, samples ranging from one single SNP up to eleven were found. In the early beginning of the outbreak in Wuhan city (between December 18-29, 2019), one to four mutations arose in the virus per patient (Shen et al. 2020), arguing that the number of fixed mutations in the world population is rapidly increasing where the infection has spread. Importantly, one Australian sample sequenced by next generation sequencing presented the USA-WA signature, suggesting that this signature is already propagated with the worldwide pandemic. Clonal mutational events within Australian samples were also observed and probably are widespread in the region. One interesting feature of the mutations in this limited number of samples is the evolutionary pressure undergone helicase gene. RNA helicases display various functions in genome replication, and it has been proposed as a therapeutic target to inhibit coronaviruses among other viruses with small molecules (Briguglio et al. 2011). Helicase evolution in SARS-CoV-2 could be rapidly occurring, making it unfeasible to drug this protein in the future. As genetic drift is allowing SARS-CoV-2 to evolve while the pandemic is still growing, the amplification efficiency of quantitative RT-qPCR tests is challenged since a single mutation even in the middle of a primer can be detrimental for PCR efficiency (Bru et al. 2008), potentially contributing to false negative results in COVID-19 testing. For this issue, we provided a way to compute the latter, by merging all variant sites called across studied samples and intersect them across primer sets available both in literature and currently in use. As new mutations can be spread depending on the founder effect, we firmly urge that primer sets for clinical testing should be tested in this way continuously, according to the current mutations found at the particular time and in the specific population which needs to be diagnosed with SARS-CoV-2 infection.

## Conclusions

We described the early mutational events in SARS-CoV-2 virus by inspecting sequencing samples from China, USA, Australia and GenBank sequenced submitted until March 31, 2020. SARS-CoV-2 variants from USA display five-point mutations with clonal (founder) pattern of spreading and considerable high allele frequencies in the next-generation sequencing samples. The latter was verified in SARS-CoV-2 sequences submitted to GenBank, since these five-point mutations displayed alleles frequencies of 50% among all USA GenBank SARS-CoV-2 sequences (n=277). SARS-CoV-2 Australian mutations were heterogeneous, but still, clonal events were found. RT-qPCR efficiency testing can be potentially affected by founder mutations, since several SNPs affects one of three primers sets currently used in COVID-19 testing.

---

**Supplementary Table 1: Detailed Characterization variants from Next Generation Sequencing (NGS) Datasets and GenBank sequences, worldwide.**

Sheet 1: Accessions and List of Mutations

Sheet 2: Variant Effect from USA NGS samples

Sheet 3: Variant Effect from Australia NGS samples

Sheet 4: Merged vcf sites

Sheet 5: GenBank variants (Worlwide)

Sheet 6: GenBank variants (USA)

## Data supplied by the author

All next generation sequencing dataset accessions and resulting computational calculations are summarized in Supplementary Table 1. All commands to obtain these computational steps are publicly available at: https://github.com/cfarkas/SARS-CoV-19_illumina_analysis. The repository can be downloaded freely.

## Competing Interest statement

José Luis Garrido is a co-founder of Ichor Biologics LLC and is currently employed by Ichor Biologics LLC, New York, United States. The rest of the authors currently works in the academia and they do not belong to industrial/comercial enterprises. All of the authors declared that they have no competing interests.

## Funding statement

Funding from the Canadian Institute of Health Research (CIHR, project grant 419220).

## Notes

https://github.com/cfarkas/SARS-CoV-19_illumina_analysis

### Competing Interest Statement

José Luis Garrido is a co-founder of Ichor Biologics LLC and is currently employed by Ichor 421 Biologics LLC, New York, United States. The rest of the authors currently works in the 422 academia and they do not belong to industrial/comercial enterprises. All of the authors declared 423 that they have no competing interests.

